# Antimicrobial susceptibility testing of *Clostridioides difficile*: a dual-site study of three different media and three therapeutic antimicrobials

**DOI:** 10.1101/2024.11.11.622946

**Authors:** Jane Freeman, Ingrid M.J.G. Sanders, Céline Harmanus, Emma V. Clark, Andrea M Berry, Wiep Klaas Smits, the European Society of Clinical Microbiology and Infectious Diseases Study Group on *Clostridioides difficile* (ESGCD)

**Author notes:** Address correspondence to Jane Freeman or Wiep Klaas Smits.

## Abstract

**Objectives:** Increasing resistance to antimicrobials used for the treatment of *Clostridioides difficile* infections necessitates reproducible antimicrobial susceptibility testing. Current guidelines take a one-size-fits-all approach and/or offer limited guidance. We investigated how the choice of medium affects measured MIC values across two sites.

**Methods:** We determined MIC values for the antimicrobials fidaxomicin, metronidazole and vancomycin for a representative collection of European *C. difficile* strains (n=235) using agar dilution on three different media: Brucella Blood Agar (BBA), Fastidious Anaerobe Agar supplemented with horseblood (FAA-HB) and Wilkins-Chalgren (WC) agar. The study was conducted at two sites to compare reproducibility. Useability (ease of preparation of the media as well as read-out of the assay) was assessed through a survey.

**Results:** We found that all media result in highly consistent aggregated MIC data for all antibiotics, with MIC_50_ and MIC_90_ within 2-fold of each other across sites. For fidaxomin, MIC values on WC were lower than on the other media. Metronidazole showed the lowest MIC on BBA, and the highest on WC. For vancomycin, there was little difference between media. Though absolute values for individual isolates differed between sites, identified resistant isolates were similar. Results obtained on FAA-HB were most consistent between sites and results obtained on WC showed the most divergence. FAA-HB was positively evaluated in the usability survey.

**Conclusions:** This study shows medium-dependent differences in *C. difficile* MICs for at least two antimicrobials across two sites. We suggest the use of FAA-HB to align with general EUCAST recommendations for susceptibility testing of anaerobic bacteria and deposited reference strains for standard susceptibility testing of *C. difficile* to increase interlaboratory reproducibility.

## INTRODUCTION

*Clostridioides difficile* is a clinically important bacterium that can cause potentially fatal gastro-intestinal disease [1]. The disease is associated with a high burden on healthcare systems, society and the economy [2-5]. At present, three antimicrobial treatments are indicated by international guidelines: fidaxomicin and vancomycin are first line treatments [6,7] except the UK, where only vancomycin is recommended as first-line)[8]. Metronidazole is no longer recommended due to clinical inefficacy unless first line therapeutics are not available or contra-indicated [6, 7]. However, despite clinical inferiority metronidazole is still widely used [8].

Reduced susceptibility or resistance of *C. difficile* has been reported for all three antimicrobials [9]. However, reported rates of resistance, particularly for metronidazole, vary widely and the reasons for this are poorly understood. Though geography and lineage of *C. difficile* might contribute to different rates of resistance, it may also attributable to differences in testing methodology [10].Current EUCAST and CLSI guidelines both advocate the use of Brucella Blood Agar for the determination of antibiotic MICs for all anaerobes, including *C. difficile*, but differ in the breakpoints defined [12, 13]

Recent studies on metronidazole resistance in *C. difficile* underscore the issues outlined above. Whereas most studies report limited metronidazole resistance [10], rates up to 44.6% have been reported in Israel [11]. Metronidazole resistance can be plasmid-mediated [12], but a subset of isolates demonstrates plasmid-independent resistance [13]. Importantly, the latter group demonstrates strong medium-dependence [13, 14]. This is suggested to be due to a mutation in the promoter of gene *nimB*, resulting in constitutive transcription [18]. Finally, it has been suggested that reduced susceptibility (MIC≥1 mg/L when tested on non-consensus medium) is associated with treatment failure [15]. Vancomycin and fidaxomicin achieve gut concentrations several orders of magnitude above the MIC of *C. difficile* isolates classified as resistant to these drugs, and therefore the clinical significance of resistance against these drugs is not yet clear. Data is lacking on whether medium composition affects vancomycin and fidaxomicin susceptibility.

Together these data clearly show the importance or routine screening of *C. difficile* using standardised conditions that might benefit from a re-evaluation of resistance breakpoints.

## METHODS

As minor differences were inevitable between sites, the catalog numbers of individual chemicals is provided as a **Supplemental Table 1**. An standard operating procedure (SOP) is available as **Supplemental Text**.

### Strain selection

We selected n=250 isolates that were collected during the COMBACTE-CDI (2018) and ClosER studies (2011-2016) [16, 17]. The isolates were selected to comprise at least 10 isolates of the top ten most prevalent European PCR ribotypes, as well as recently emerging ribotypes or ribotypes associated with multi-drug resistance (resistance to>3 antimicrobial classes) (**Supplemental Table 2**). We included isolates that were previously identified as showing reduced susceptibility towards metronidazole (n=39), vancomycin (n=31) or fidaxomicin (n=1) [16, 17].

To reduce differences arising from repeated sub-culturing, *C. difficile* isolates were subcultured on cycloserine-cefoxitin egg yolk agar (E&O laboratories, Bonnybridge, Scotland, UK) for 48h in Leeds. Duplicate sets of glycerol stocks were prepared by resuspending the growth in glycerol broths, dividing into two aliquots and freezing at -80°C; one aliquot was sent on dry ice to Leiden. Control strains (*C. difficile* ATCC 700057, *C. difficile* E4 [18], *Bacteroides fragilis* ATCC 25285 and *Enterococcus faecalis* ATCC 29212) were similarly shared between both sites.

### Antimicrobial susceptibility testing

*C. difficile* isolates were removed from -80°C storage and subcultured anaerobically to ensure purity, prior to inoculation of pre-reduced Schaedler’s anaerobic broth for 24h [13]. Isolates were transferred to pre-reduced sterile saline and adjusted to McFarland standard 1.0. Non-antibiotic containing plates were incubated aerobically and anaerobically.

Antibiotic-containing agar plates were prepared by mixing 2mL dilution of the antimicrobial with 18 mL molten agar and distributing into petri dishes. Blood, haemin and Vitamin K were added after autoclaving and prior to distribution into petri dishes (**Supplemental Text**). Fidaxomicin was dissolved in 100% DMSO as a solvent, and further diluted into 10% DMSO as a diluent.

Metronidazole was dissolved in 100% DMSO as a solvent, and further diluted into water as a diluent. Vancomycin was dissolved in water as a solvent, and further diluted into water as a diluent. The final concentrations of the antimicrobials in the agar dilution experiments were 0.015-16 mg/L for fidaxomicin, 0.125-32 mg/L for metronidazole and 0.125-32 mg/L for vancomycin.

Saline suspensions of *C. difficile* isolates were inoculated onto agar plates using a multipoint inoculator and incubated anaerobically for 48h. The minimum inhibitor concentration is defined as the lowest dilution at which growth is completely inhibited or where only a single colony remains.

### Analysis and visualisation

Plates were read by two technicians and results were logged only when control strains demonstrated MIC values within a predefined range (**Table 1**). For fidaxomicin no breakpoints were defined at the time of testing (2019). We therefore defined breakpoints for this study as ≤1 mg/L: susceptible; >1 mg/L: resistant, in line with those used in the CLosER study [16]. For metronidazole and vancomycin, EUCAST breakpoints at the time of testing were used (≤2 mg/L: susceptible, >2 mg/L: resistant. MIC_50_ and MIC_90_ were determined on the basis of ranked MIC values.

**Table 1.**
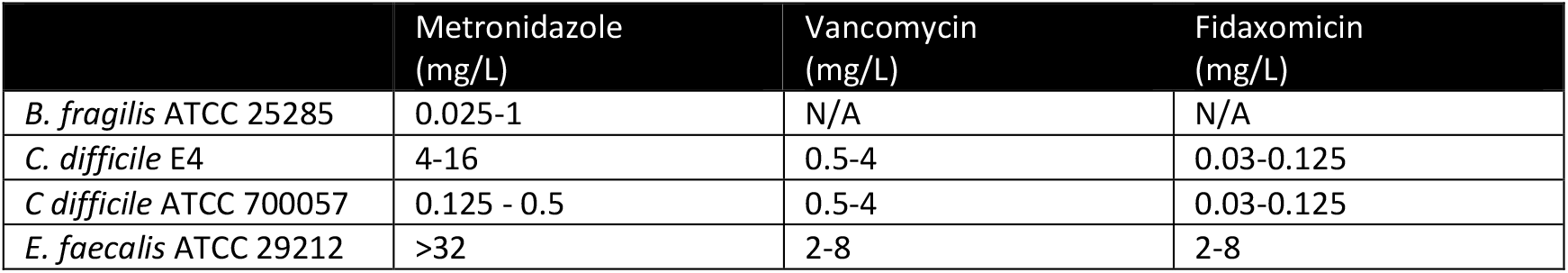
Expected MIC ranges for control strains.

Data were collected in Microsoft Excel and converted into a tidy format (**Supplemental Table 2**). Graphs were generated in SuperPlotsOfData [19] and Eulerr [20], and further compiled in Adobe Illustrator 2022 (26.3.1).

## RESULTS

### Agar dilution provides highly similar overall susceptibility data between sites

Of the n=250 strains that were initially selected, some could not be revived from the original stocks. As a result, n=235 isolates were shared between laboratories in Leeds and Leiden. During the experiments in the Leeds laboratory, 3 additional isolates we judged not to contain *C. difficile* upon subculture and were excluded from the subsequent analysis.

MIC_50_ and MIC_90_ values were generally within 2-fold of each other between sites; only for the vancomycin MIC_50_, a 4-fold difference was observed on BBA medium. When differences were observed, MIC values were higher at the LUMC site in comparison to the Leeds site. This suggests that subtle differences in experimental procedures may lead to a systematic difference in MIC values measured.

MIC ranges show a similar trend; when differences are observed, these are mostly at the lower end of the MIC range and maximum MIC values are within 2-fold of each other. Any differences are therefore unlikely to affect the qualification of strains as resistant.

### Fidaxomicin and metronidazole show medium-dependent differences in antimicrobial susceptibility

Next, we compared the distribution in MIC values observed for each combination of medium and antibiotic. Overall, we observed a strikingly similar pattern for values determined in Leeds (**Figure 1A**) and Leiden (**Figure 1B**).

**Figure 1.**
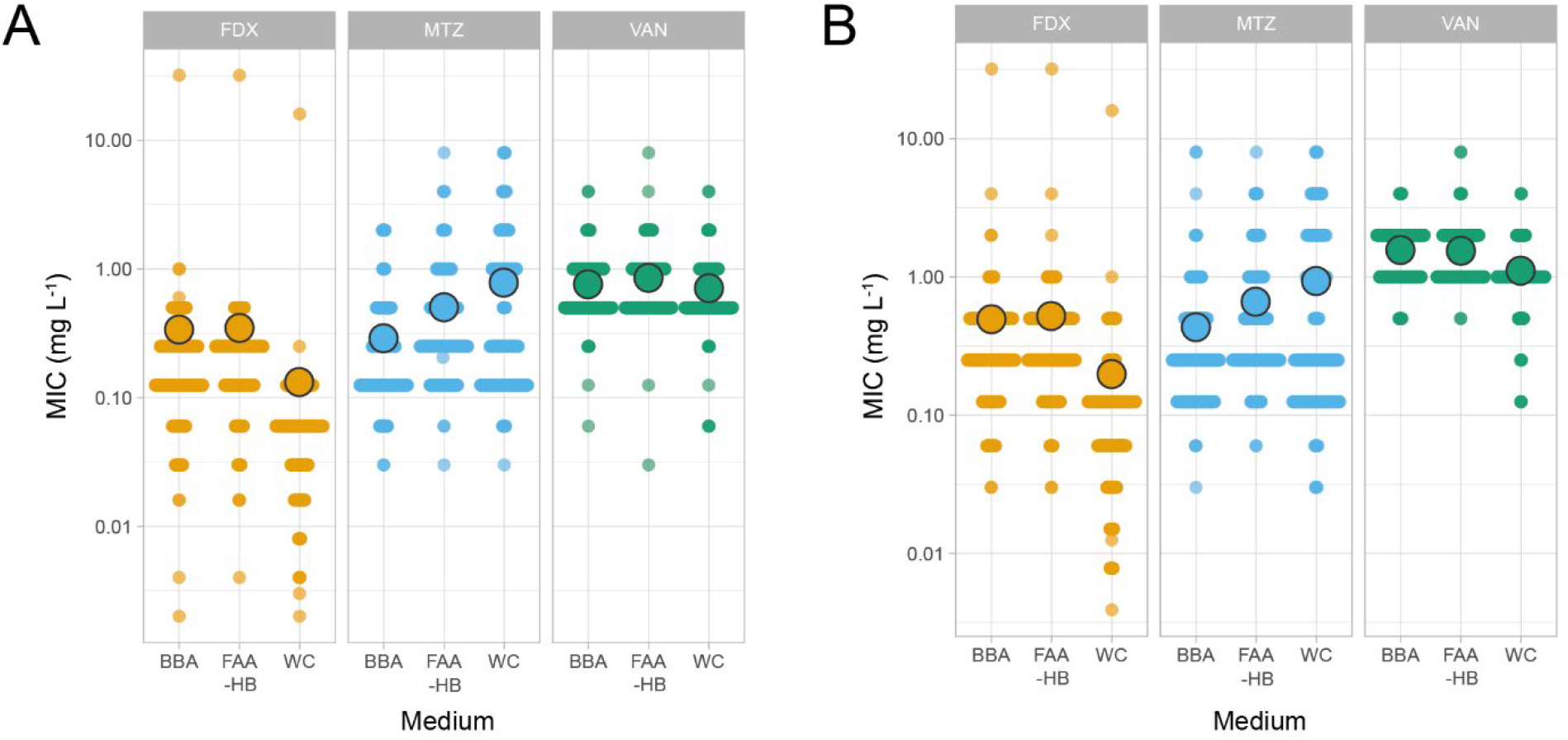
Comparison of MIC values for Leeds (A) and Leiden (B). Individual datapoints are shown, with the mean indicated with the largest circle. Data is shown for fidaxomicin (FDX, ochre), metronidazole (MTZ, blue) and vancomycin (VAN, green) on three different media: brucella blood agar (BBA), fastidious anaerobe agar (FAA-HB) and Wilkins-Chalgren (WC). For details see Methods.

As also noted above, mean values determined at Leiden appear to be slightly higher than those determined in Leeds. However, within sites, the different antimicrobial-medium combinations show similar trends. For fidaxomicin, we find that mean MICs are similar for FDX and MTZ, but are markedly lower for WC medium. This suggest that the use of WC in agar dilution experiments may lead to a systematic underestimation of potential reduced susceptible isolates, compared to other media. An opposite pattern is observed for metronidazole: mean MIC values progressively increase from BBA, through FAA-HB to WC medium. Thus, for both fidaxomicin and metronidazole there is a strong medium-dependent effect on susceptibility. For vancomycin, results indicate a lesser medium-dependence with mean values being similar for all media in the Leeds dataset, and only WC resulting in slightly lower mean MIC values in the Leiden dataset.

### Interlaboratory differences differ per medium used

Though aggregated data shows highly similar trends (**Table 2, Figure 1**), this analysis could potentially mask differences between the MIC values for individual isolates obtained in the two laboratories. In order to assess this, we calculated the ratio of the MIC values for each isolate; if data were 100% congruent, this should result in a ratio of 1 (the same MIC value). We find that the mean value indeed approaches this (**Figure 2A**), in particular for FDX on all media.

**Table 2.** Aggregated MIC values. MIC values for individual isolates were aggregated to the concentration that inhibits the growth of >50% of all isolates (MIC50) or >90% of all isolates (MIC90). Data is shown for fidaxomicin (FDX), metronidazole (MTZ) and vancomycin (VAN) on three different media: brucella blood agar (BBA), fastidious anaerobe agar (FAA-HB) and Wilkins-Chalgren (WC). For details see Methods.

| Site | Antimicrobial | Medium | MIC<sub>50</sub><br>(mg/L) | MIC<sub>90</sub><br>(mg/L) | MIC range (mg/L) |
| --- | --- | --- | --- | --- | --- |
| Leeds | FDX | BBA | 0.125 | 0.5 | 0.002-32 |
|  |  | FAA-HB | 0.25 | 0.5 | 0.004-32 |
|  |  | WC | 0.06 | 0.125 | 0.002-16 |
|  | MTZ | BBA | 0.25 | 0.5 | 0.03-2 |
|  |  | FAA-HB | 0.25 | 1 | 0.03-8 |
|  |  | WC | 0.25 | 2 | 0.03-8 |
|  | VAN | BBA | 0.5 | 1 | 0.06-4 |
|  |  | FAA-HB | 0.5 | 1 | 0.03-8 |
|  |  | WC | 0.5 | 1 | 0.06-4 |
| Leiden | FDX | BBA | 0.25 | 0.5 | 0.03-32 |
|  |  | FAA-HB | 0.25 | 0.5 | 0.03-32 |
|  |  | WC | 0.125 | 0.25 | 0.004-16 |
|  | MTZ | BBA | 0.25 | 1 | 0.03-8 |
|  |  | FAA-HB | 0.25 | 2 | 0.06-8 |
|  |  | WC | 0.25 | 2 | 0.03-8 |
|  | VAN | BBA | 2 | 2 | 0.5-4 |
|  |  | FAA-HB | 1 | 2 | 0.5-8 |
|  |  | WC | 1 | 2 | 0.125-4 |

**Figure 2.**
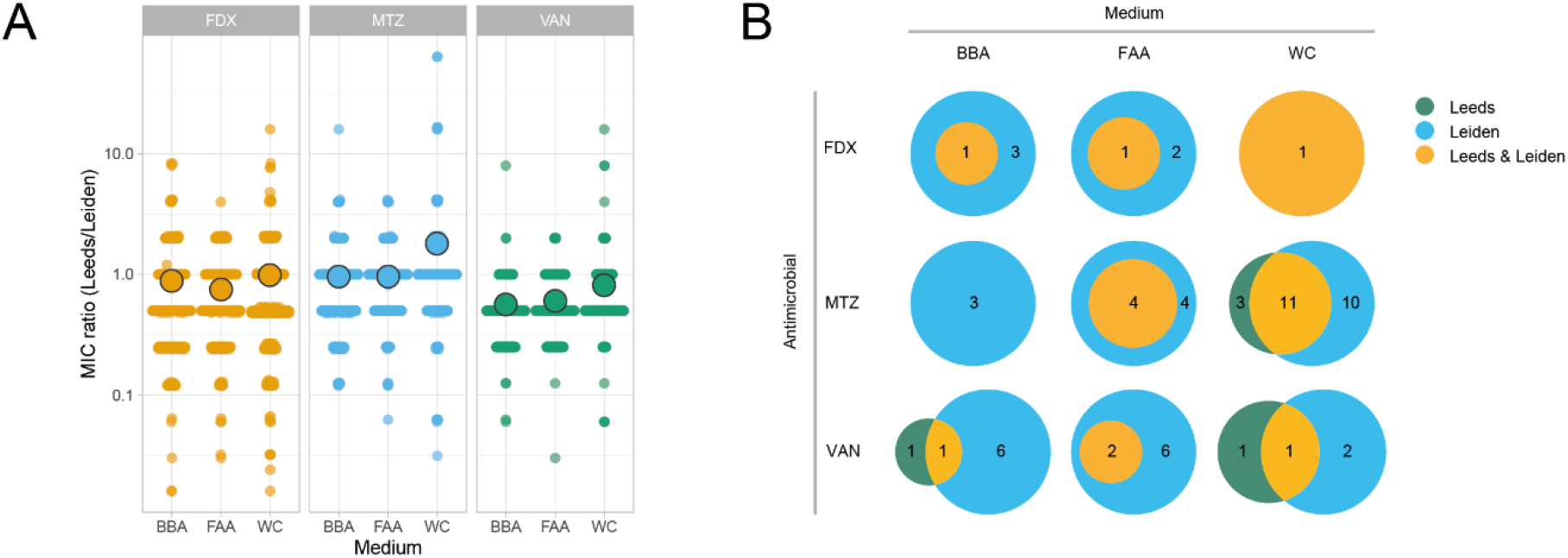
Inter-site reproducibility of antimicrobial susceptibility testing. **A**. Ratios of MIC values obtained at the two sites were calculated for each isolate. Individual datapoints are shown, with the mean indicated with the largest circle. Data is shown for fidaxomicin (FDX, ochre), metronidazole (MTZ, blue) and vancomycin (VAN, green) on three different media: brucella blood agar (BBA), fastidious anaerobe agar (FAA-HB) and Wilkins-Chalgren (WC). **B**. The number of isolates with MIC≥1 mg/L (FDX) or MIC≥4 mg/L (MTZ and VAN) under one or more condition were compared. A value of 1 reflects indicates that 1 isolate met this criterium on one medium. For details see Methods.

**Figure 3.**
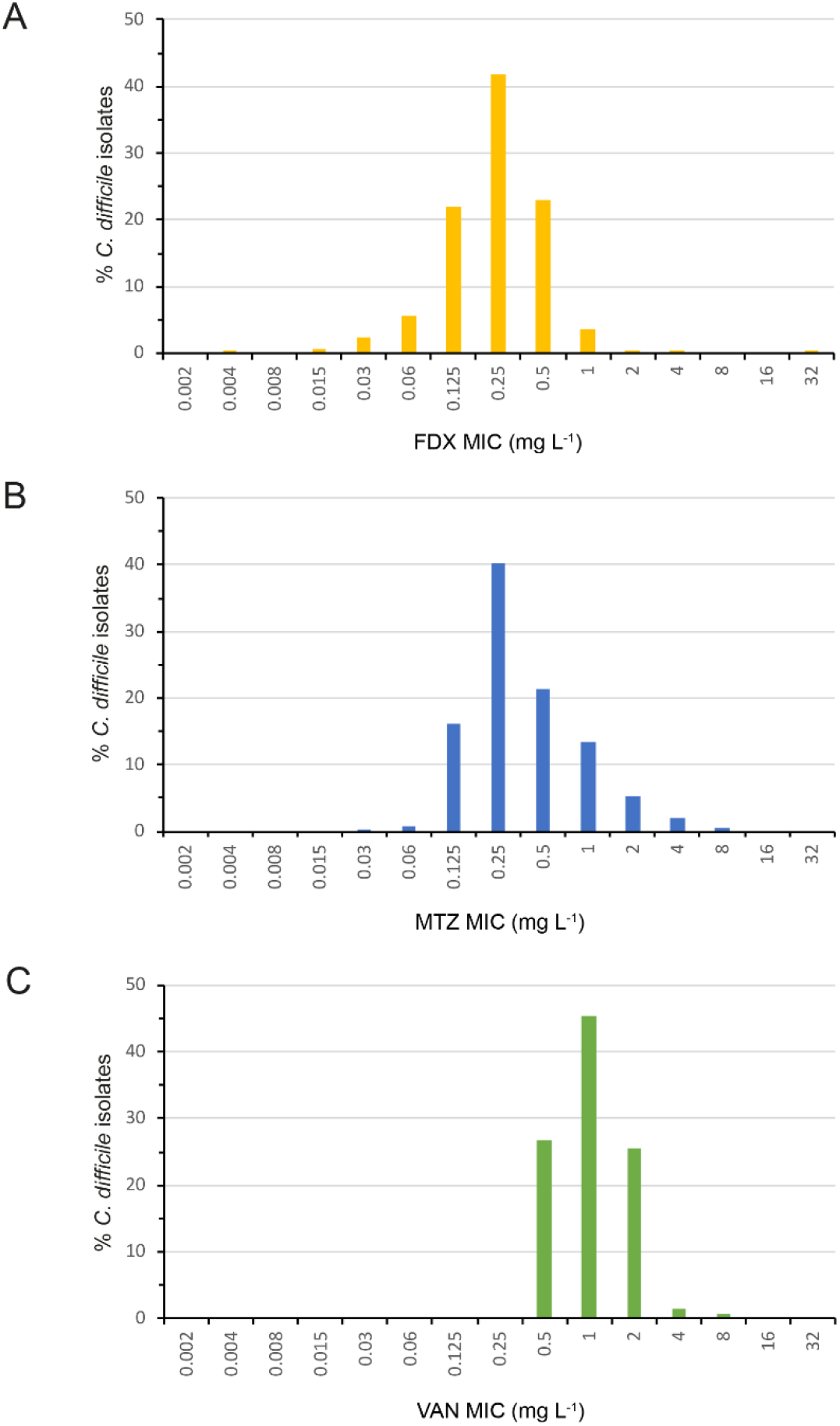
MIC distributions. Aggregated data for MICs determined by FAA-HB agar dilution method across both sites for fidaxomicin (orange) metronidazole (blue) and vancomycin (green).

A high reproducibility should result in a tight clustering of datapoints. We note that data obtained on FAA-HB medium shows a narrower distribution of ratios, compared to BAA and WC (**Figure 1A**).

Next, we assessed how medium-antimicrobial combinations affect the identification of resistant isolates, which we deem to be the clinically most relevant outcome parameter. We identified all isolates with an MIC > 1 mg/L (FDX) or MIC>2 mg/L (MTZ or VAN) (**Supplemental Table 2**) on one or more medium-antimicrobial combinations. As expected, the number of isolates identified in this way was higher for the Leiden site than the Leeds site, consistent with the higher mean MIC values.

However, in general, those isolates identified in Leeds were also identified in Leiden (**Figure 2B**). Notably, however, identifications of vancomycin resistant isolates and/or identifications on WC medium showed a significant laboratory-specific effect. The most consistent results were observed for FAA-HB medium, where all isolates identified as resistant in Leeds were also identified in Leiden.

### Distinct media offer easier handling or read-out

We considered several aspects that might contribute to experimental variability in our data. In particular steps necessary to prepare media, as well as the ability to clearly read the MIC data stemming from the agar dilution data appeared to be important sources of variation to us. We therefore queried the teams that performed the experiments about these aspects. Four members of staff were surveyed, with three giving feedback using Likert scales on ease of preparation, readability and detection of contamination (**Supplemental Table S3**). The currently CLSI-recommended Brucella Blood Agar consistently scored least well on all survey questions and was the least preferred option among staff performing the procedure.

## DISCUSSION

Here, we determined how the choice of medium affects antimicrobial susceptibility testing for *C. difficile* for therapeutic antimicrobials fidaxomicin, metronidazole and vancomycin. We find that agar dilution offers a reliable assessment of susceptibility with minor systematic differences between sites using a harmonized protocol. There were mechanical differences in agar preparation method between the two sites (Leeds used an automated agar preparation, while Leiden used a hand-poured agar technique) which may account for such minor differences. Nonetheless, we found clear evidence for medium-dependent susceptibility for fidaxomicin and metronidazole, and show that determining susceptibility on FAA-HB medium shows the highest inter-laboratory reproducibility. FAA-HB also scored highly for ease of use, ease of reading MICs and detection of contamination among staff performing the procedure, in contrast to BBA, which was consistently the lowest scoring medium.

Medium dependent resistance in *C. difficile* has so far only been described for metronidazole [13, 14], and medium-dependent effects on fidaxomicin susceptibility have to the best of our knowledge not been documented before. Though the use of WC could clearly result in lower MIC values for fidaxomicin (and possibly vancomycin), it should be noted that this does not affect the identification of highly resistant strains [21]. FDX-reduced susceptible strains have rarely been identified but the inclusion of the only FDX-resistant *C. difficile* isolate so far identified in large scale surveillance studies [20,24] gives confidence that FDX resistance would be detected on the recommended FAA-HB medium. For metronidazole, heme in the medium appears to be a key determinant for resistance, and it should be noted that when using any fresh blood based medium for determining metronidazole MICs in *C. difficile*, blood should be fresh, to avoid degradation of haemin [13, 14].

The mechanisms behind metronidazole resistance are only recently being elucidated: plasmid-mediated and *nim* gene expression. Importantly, the use of FAA-HB as recommended on the basis of this study can detect both types of metronidazole resistance. The reason for increased susceptibility for fidaxomicin (and potentially vancomycin) on WC medium is at present unknown, but may be influenced by the more defined and minimal nature of WC, compared to the blood-supplemented media (FAA-HB and BBA). This may marginally increase growth and accentuate differences at the lower end of concentration ranges seen for fidaxomicin.

The strengths of the present study include the well-controlled setup and the inclusion of a large number of representative *C. difficile* isolates. Our data is limited by the fact that we did not perform an in-depth analysis of potentially discrepant results, and did not perform repeat testing of the same set of isolates.

Our study suggests that the use of BBA medium for agar dilution as recommended in some current susceptibility testing guidelines (e.g. CLSI) might lead to an underestimation of in particular metronidazole resistance and could contribute to inter-laboratory variation in reported MIC values. Our data support the use of FAA-HB medium for susceptibility testing in *C. difficile*, since it offers a good balance of usability, reproducibility and sensitivity. Notably, this is in line with studies of disk susceptibility in other anaerobes, including *Bacteroides*/*Phocaeicola*/*Parabacteroides, Prevotella, Fusobacterium, Clostridium* and *Cutibacterium* and now part of EUCAST recommendations [22, 23].

The increasing number of reports of *C. difficile* with reduced susceptibility or resistance to the antibiotics tested here [12, 21, 24-26] warrants a systematic surveillance of drug resistance for this organism and a re-evaluation of breakpoints. At present, prevalence of resistant isolates varies strongly between reports and is subject of dispute [26, 27]. A consensus method for susceptibility testing can contribute to inter-laboratory reproducibility. These data support the adoption of FAA-HB as the optimal medium for determining *C. difficile* susceptibility to relevant agents for treatment. They also support the current EUCAST breakpoints for metronidazole, vancomycin and fidaxomicin (≤ 2mg/L: susceptible, >2 mg/L: resistant; v14.0, 01-Jan-2024)[28], while noting that the panel were selected to include isolates previously showing reduced susceptibility to metronidazole and fidaxomicin. The panel included only one *C. difficile* isolate known to harbour fidaxomicin resistance (16 mg/L), with the remaining isolates demonstrating susceptibility 0.004 and 4mg/L. More than 99% of *C. difficile* isolates were susceptible at <1mg/L fidaxomicin, and >95% at <0.5mg/L on FAA-HB (aggregated data from both sites). To further assist laboratories undertaking susceptibility testing of *C. difficile* isolates, we have deposited isolates that were identified as resistant against FDX, MTZ and VAN in both laboratories at the NCTC for use as reference strains by other laboratories (**Table 3**) in addition to the guideline-recommended *C. difficile* ATCC 70057. Use of these strains in future work will facilitate interlaboratory comparisons of absolute MIC values.

**Table 3.** Recommended *C. difficile* strains for antimicrobial susceptibility testing of *C. difficile* on FAA-HB medium with MIC values as determined in this study.

| Isolate | NCTC accession | Fidaxomicin MIC (mg/L) | Metronidazole MIC (mg/L) | Vancomycin MIC (mg/L) | Description | Reference |
| --- | --- | --- | --- | --- | --- | --- |
| 15-7365627 | NCTC 15114 | 0.25 | 4 | 0.5-2 | RT016. Medium dependent resistance#. | This study |
| E4 | NCTC 15085 | 0.125 | 4-16 | 0.5-1 | RT010 | [30] |
| L_16.7570132 | NCTC 15086 | 32 | 0.25 | 1.0 | RT344 | [21], this study |
| L_13.7933412 | NCTC 15087 | 0.125-0.25 | 0.125-0.25 | 8 | RT356 | This study |

The results from the present study have been communicated to EUCAST and will be recommended for determination of *C. difficile* MICs by agar dilutions in the next revision of their guidance.

Beyond the relevance for the epidemiological monitoring of the emergence of strains resistant to treatment antimicrobials, the approach proposed here may help to effectively guide healthcare professionals in treating CDI patients.

## Transparency declaration

### Conflict of interest

JF has received research funding from Pfizer and Crestone, and speaking honoraria and sponsorship to attend meetings from Tillotts.

WKS has performed research for Cubist Pharmaceuticals, and holds a public-private partnership grant with Acurx Pharmaceuticals.

All other authors: none to declare.

## Acknowledgements

The authors wish to acknowledge members of the Leeds and Leiden research groups for discussions.

## Access to data

All data is contained in the manuscript and the provided Supplemental Materials.

## Contribution

JF: writing, supervision, writing-original draft; CH: investigation; IMJGS: investigation; EVC: investigation; AMB investigation; WKS: formal analysis, data curation, writing-original draft, visualization, supervision. All authors reviewed and edited the manuscript, and approved the final version.

