## Supplementary material for "Antimicrobial susceptibility testing of *Clostridioides difficile*: a dual-site study of three different media and three therapeutic antimicrobials": Supplemental Table S1.docx

**Supplement S1. Reagents and catalogue numbers**

Leeds site:

| **Reagent** | **Supplier** | **Catalogue number** |
| --- | --- | --- |
| Pre-poured Brazier’s CCEY agar plates | E&O laboratories | PP4070, |
| Columbia Agar & 5% Horse Blood | E&O laboratories | PP0120, |
| Schaedlers anaerobic broths | Oxoid/Thermofisher | CM0497B |
| Saline 0.9% | E&O labs | BM0381 |
| Wilkins Chalgren Agar base | Oxoid/Thermofisher | CM0619 |
| Brucella Agar base | Oxoid/Thermofisher | CM0169 |
| Laked Sheeps Blood | E&O labs | DSC |
| Fastidious anaerobe agar base | LabM | LabM090 |
| Defibrinated Horse Blood | E&O laboratories | DHB |
| Metronidazole | Sigma | M3761 |
| Vancomycin | VWR | Applichem A1839 |
| Fidaxomicin | Stratech | Selleckchem S4277-SEL |
| Haemin | Sigma | 1003093408 |
| Vitamin K | Thermo Scientific | L10575.06 |

Leiden site:

| **Reagent** | **Supplier** | **Catalogue number** |
| --- | --- | --- |
| Tryptic Soy Sheep Blood Agar plates (TSS) | bioMérieux | 43009 |
| Schaedlers anaerobic broth | Oxoid/Thermofisher | CM0497B |
| Saline 0.9% | In-house; prepared with Sodium chloride, Supelco | 1.06404.1000 |
| Wilkins Chalgren Agar base | Oxoid/Thermofisher | CM0619 |
| Brucella Agar base | Oxoid/Thermofisher | CM0169 |
| Sheep blood, defibrinated | Xebios Diagnostics GmbH | 10000100/10000250 |
| Fastidious anaerobe agar base | Biotrading Benelux BV | Neogen NCM0014a |
| Horse Blood, defibrinated | Xebios Diagnostics GmbH | 2000100 |
| Metronidazole | Sigma | M3761 |
| Vancomycin Hydrochloride | VWR | Applichem A1839 |
| Fidaxomicin | Bioconnect BV | Selleckchem S4277 |
| Hemin | Sigma | 51280 |
| Vitamin K1 | Carl Roth GmbH & Co | 3804.2 |
