## Supplementary material for "Antimicrobial susceptibility testing of *Clostridioides difficile*: a dual-site study of three different media and three therapeutic antimicrobials": Supplemental Table S3.docx

**Supplemental Table S3.** Survey of staff experience of using different agars to perform susceptibility testing for MICs

| **1. How did you prepare your agar?** | preparator | preparator | hand pour |  | Average | Comments | | | | |
| --- | --- | --- | --- | --- | --- | --- | --- | --- | --- | --- |
| **2. How easy did you find preparing BBA?** Difficult \| \| \| \| neither difficult nor easy \| \| \| \| \| Very easy; 1 \| 2 \| 3 \| 4 \| 5 \| 6 \| 7 \| 8 \| 9 \| 10 | 9 | 3 | 8 |  | 6.7 | The media was easy to prepare, issues were with plate pourer | | 4 reagents required, so takes longer to prepare. Sheeps blood must also be laked which requires further steps (aliquot, freeze, thaw) | |  |
| **3, How easy did you find preparing FAA?** Difficult \| \| \| \| neither difficult nor easy \| \| \| \| \| Very easy; 1 \| 2 \| 3 \| 4 \| 5 \| 6 \| 7 \| 8 \| 9 \| 10 | 9 | 10 | 8 |  | 9.0 | The media was easy to prepare, issue were plate pourer problems | | Only agar and DHB required | |  |
| **4. How easy did you find preparing WC?** Difficult \| \| \| \| neither difficult nor easy \| \| \| \| \| Very easy; 1 \| 2 \| 3 \| 4 \| 5 \| 6 \| 7 \| 8 \| 9 \| 10 | 10 | 10 | 10 |  | 10.0 | The media was easy to prepare, issue were plate pourer problems | | Only agar and haemin required | | No blood |
| **5. Which agar took the longest to prepare (please rank 1,2 or 3)** | | | | | | | | | | |
| **BBA** | 1 | 1 | 1 |  |  | More likely to make mistakes as there are additions to make after autoclaving but before pouring | | There are more reagents to add which allows for a greater chance of error | | No errors, basic stuff to prepare |
| **FAA** | 2 | 2 | 2 |  |  | More likely to make mistakes as there are additions to make after autoclaving but before pouring | | I *think* I forgot to add blood once | | No errors, basic stuff to prepare |
| **WC** | 3 | 3 | 3 |  |  |  | | This is our standard media type, so errors didn’t occur – haemin is added before autoclave so you are unlikely to forget | | No errors, basic stuff to prepare |
| **6. How often do you feel errors were made in preparing BBA** Often \| \| \| \| Sometimes \| \| \| \| \| Rarely; 1 \| 2 \| 3 \| 4 \| 5 \| 6 \| 7 \| 8 \| 9 \| 10 | 8 | 7 | 10 |  | 8.3 | The dark colour of the agar made it ifficult to read when growth was light. Also harder to see the innuclation point marks in the agar. | | C. diff growth blends into the colour of the agar – extremely difficult to determine endpoint results or when growth becomes +/- | | all plates where easy to read |
| **7. How often do you feel errors were made preparing FAA** Often \| \| \| \| Sometimes \| \| \| \| \| Rarely; 1 \| 2 \| 3 \| 4 \| 5 \| 6 \| 7 \| 8 \| 9 \| 10 | 8 | 9 | 10 |  | 9.0 | The dark colour of the agar made it ifficult to read when growth was light. Also harder to see the innoculation point marks in the agar. | | C.diff growth looks great, the only downside is you can’t overlay the plates as the agar is opaque | |  |
| **8. how often do you feel errors were made preparing WC** Often \| \| \| \| Sometimes \| \| \| \| \| Rarely; 1 \| 2 \| 3 \| 4 \| 5 \| 6 \| 7 \| 8 \| 9 \| 10 | 10 | 10 | 10 |  | 10.0 |  | | C.diff growth looks great and you can overlay the plates to read, if this is your preferred method | |  |
| **9. How easy did you find reading MIC endpoints on BBA** Difficult \| \| \| \| neither difficult nor easy \| \| \| \| \| Very easy; 1 \| 2 \| 3 \| 4 \| 5 \| 6 \| 7 \| 8 \| 9 \| 10 | 4 | 1 | 8 |  | 4.3 | Again darker aggr made it difficult to see innoculatio points making it hard to see if the contaminating growth was where the innoculation was made | | Colour of agar can make this tricky | |  |
| **10. How easy did you find reading MIC endpoints on FAA** Difficult \| \| \| \| neither difficult nor easy \| \| \| \| \| Very easy; 1 \| 2 \| 3 \| 4 \| 5 \| 6 \| 7 \| 8 \| 9 \| 10 | 4 | 9 | 8 |  | 7.0 | Again darker agar made it difficult to see innoculation points making it hard to see if the contaminated growth was where the innolcuation was made | | Contaminants contrast the colour of the DHB nicely | |  |
| **11. How easy did you find reading MIC endpoints on WC** Difficult \| \| \| \| neither difficult nor easy \| \| \| \| \| Very easy; 1 \| 2 \| 3 \| 4 \| 5 \| 6 \| 7 \| 8 \| 9 \| 10 | 9 | 10 | 8 |  | 9.0 |  | | Routine agar type - trained eye to spot contaminants | |  |
| **12. How easy did you find it to detect contamination (non- *C.difficile* bacteria) on BBA** Difficult \| \| \| \| neither difficult nor easy \| \| \| \| \| Very easy; 1 \| 2 \| 3 \| 4 \| 5 \| 6 \| 7 \| 8 \| 9 \| 10 | 4 | 3 | 8 |  | 5.0 |  | | | | |
| **13. How easy did you find it to detect contamination (non- *C.difficile* bacteria) on FAA** Difficult \| \| \| \| neither difficult nor easy \| \| \| \| \| Very easy; 1 \| 2 \| 3 \| 4 \| 5 \| 6 \| 7 \| 8 \| 9 \| 10 | 4 | 9 | 8 |  | 7.0 |  |  |  |  |  |
| **14. How easy did you find it to detect contamination (non- *C.difficile* bacteria) on WC** Difficult \| \| \| \| neither difficult nor easy \| \| \| \| \| Very easy; 1 \| 2 \| 3 \| 4 \| 5 \| 6 \| 7 \| 8 \| 9 \| 10 | 8 | 10 | 5 |  | 7.7 |  |  |  |  |  |
| **Which agar would you prefer to work with when determining C. difficile MICs? Please rank 1,2 or 3** |  |  |  |  |  |  |  |  |  |  |
| **BBA** | 3 | 3 | 3 |  | 3.0 |  |  |  |  |  |
| **FAA** | 2 | 2 | 1 |  | 1.7 |  |  |  |  |  |
| **WC** | 1 | 1 | 2 |  | 1.3 |  |  |  |  |  |
