## Supplementary material for "Antimicrobial susceptibility testing of *Clostridioides difficile*: a dual-site study of three different media and three therapeutic antimicrobials": Supplemental Text.docx

### Simplified protocol for agar dilution of *C. difficile* strains on FAA-HB

This protocol is meant as a guide for setting up susceptibility testing of *C. difficile*. It is not intended as a standard operating procedure, and we recommend laboratories use this document to set up their own SOPs detailing use of dedicated equipment with relevant instructions.

#### Culturing from storage

1. Recover pure *C. difficile* isolate on a solid agar medium (CCEY, TSS or similar); incubate for 48h at 37⁰C under anaerobic conditions.
2. Label a tube with 4 mL Schaedler’s anaerobic broth and a tube with 4mL 0.9% saline per strain, and prereduce for a minimum of 24h at 37⁰C under anaerobic conditions. Include extra tubes that will not be inoculated and serve as negative controls.
3. Inoculate the strains into the prereduced Schaedler’s anaerobic broth, mock inoculate the negative control broth.
4. Note that for non-*C. difficile* control strains, incubation times may be shorter as no pre-culture is necessary (see section Inoculating the plates). Adjust based on strains to be included.

#### Preparation of diluent

1. For metronidazole and vancomycin: diluent is sterile water.
2. For fidaxomicin: diluent is 10% sterile DMSO.
3. Prepare and label 50 mL centrifuge tubes according to Table 1 (note: for metronidazole and vancomycin it is generally enough to assess the MIC down to 0.03 mg L^-1^, for fidaxomicin it is recommended to go down to 0.002 mg L^-1^)

Table 1 Labelling and volumes for diluent tubes

| Volume of diluent | 20mL | 30mL | 42mL |
| --- | --- | --- | --- |
| Labels | 32 | 16 | 8 (B) |
|  | 4 | 2 | 1 (C) |
|  | 0.5 | 0.25 | 0.125 (D) |
|  | 0.06 | 0.03 | 0.015 (E) |
|  | 0.008 | 0.004 | 0.002 |

#### Preparation of antimicrobial-containing plates

1. Label duplicate petri dishes with the antimicrobial and the concentration (e.g. M 32 A, M 16 A etc, M 32 B, M16 B etc).
2. Prepare stocks for the antimicrobials according to Table 2 (label: STOCK)
3. Prepare working stock solutions (A) of 640 mg L^-1^ for the antimicrobials by diluting the stock solution 1:10 in diluent
4. Add 20mL of A to tube “32”, 10 mL of A to tube “16” and 6mL of A to tube “8 (B)”
5. Add 20ml of B to tube “4”, 10mL of B to tube “2”, and 6mL of B to tube “1 (C)”
6. Continue this until all dilutions are made.
7. Add 2 mL of each solution to the corresponding empty, labelled, petri dishes; for the plates without antimicrobials (“0”) add 2mL of diluent only (prepare 2 plates without antibiotics, “0+O_2_” will be incubated aerobically and “0” will be incubated anaerobically).
8. Prepare sufficient amount of agar to allow for all plates to be poured, see Table 3.
9. Dry plates (e.g. 20 minutes in a 37C)

Table 2 Preparation of stock and working stock solution of antimicrobials

| Antimicrobial | Minimal MIC range to test (mg L^-1^) | Solvent | Diluent | Stock concentration | Amount of powder (mg) | Volume of solvent (mL) |
| --- | --- | --- | --- | --- | --- | --- |
| Vancomycin | 0.03-32 | Water | Water | 6400 mg L^-1^ | 32 | 5 |
| Metronidazole | 0.03-32 | DMSO | Water | 6400 mg L^-1^ | 32 | 5 |
| Fidaxomicin | 0.002-32 | DMSO | 10% DMSO | 6400 mg L^-1^ | 32 | 5 |

Table 3 Preparation of agar medium

| Number of plates | 1 | 50 | 200 |
| --- | --- | --- | --- |
| Fastidious Anaerobe Agar base (g) | 0.92 | 46 | 184 |
| Water | 17 | 850 | 3400 |
| Autoclave, let cool to hand warm | | | |
| Horse blood  (add to medium after autoclaving) | 1 | 50 | 200 |
| Dispense 18 mL of FAA-HB agar to petridish containing antimicrobial and mix well | | | |
| Antimicrobial or diluent | 2 | 100 | 400 |
| Total volume (mL) | 20 | 1000 | 4000 |

#### Inoculating the plates

1. Remove the anaerobic broths from incubator, and into a sterile laminar flow cabinet, along with the corresponding sets of 0.9% saline. Only proceed if the blank (non-inoculated) culture is clear.
2. Dilute cultures into saline to achieve McFarland Standard 1.0 (approximately 1:10, adjust as necessary)
3. Note: for non-*C. difficile* control strains – use a swab/loop to inoculate sterile saline to achieve a uniform McFarland Standard 1.0 solution
4. Ideally, use a multipoint inoculator (according to instructions of the manufacturer) to inoculate each set of plates, starting from the “0” plates and progressing to the “32” plate. If no multipoint inoculator is available, manually spot 5 μL of the McFarland standard 1.0 solution per strain. Make sure to log the position of each strain and mark the positioning of the multipoint inoculator.
5. Leave the plates to dry before stacking the plates upside down and incubating them anaerobically at 37⁰C. Incubate the “0+O_2_” plate (without antimicrobial) aerobically at 37⁰C.
6. For a new set of plates, switch pins/pin head on the multipoint inoculator, or sterilize.

#### Reading the MICs

1. MICs should be read out after 48h incubation at 37⁰C under anaerobic conditions. Optionally, and additionally, MICs can also be read out at 24h.
2. Record only data for *C. difficile* strains that show growth on the anaerobic “0” plate; if no growth is observed on this plate, the test for this strain should be repeated. The result should be documented.
3. If the aerobically incubated “0 +O_2_” plate shows growth for *C. difficile*, the test for this strain should be repeated. Contamination should be noted on the result form.
4. Duplicate plates should show no more than 1 dilution difference in MIC. If larger, repeat the experiment.
5. Interpret data only if control strains included in the experiments show MICs within the expected range (dependent on the strains used). If control MICs are out of range, repeat the experiment. Note on the file that control MICs were out of range.
6. Record the results according to Table 4. Note down growth (+), approximate 50% inhibition of growth (+/-), a single colony (1) and no growth (NG). The MIC is the highest dilution showing 1 colony or NG. Ideally, record images of each individual plate as raw data.
7. Results should be independently read by two scientists; results are reviewed by the Project lead and documented in the Project file.

Table 4 Reporting table for agar dilution results

| MIC t=48h | Plate | | | | |  |
| --- | --- | --- | --- | --- | --- | --- |
| Strain | Abx  0+O^2^ | Abx  0 | Abx  0.002 | …  [insert relevant number of columns] | Abx  32 | Comment |
| 1 A |  |  |  |  |  |  |
| 1 B |  |  |  |  |  |  |
| 2 A |  |  |  |  |  |  |
| 2 B |  |  |  |  |  |  |
| ….  [insert relevant number of strains] |  |  |  |  |  |  |

Key: + = growth, +/- = approximate 50% inhibition, 1 = single colony, - = no growth.
